# Bayesian parameter estimation in non-stationary semiflexible polymers from ensembles of trajectories

**DOI:** 10.1101/484691

**Authors:** Christopher A. Penfold

## Abstract

During the cell-cycle and meiosis, during development, or in response to stress, chromosomes undertake dramatic programs of reorganisation, which can result in major changes to genomic architecture, as well as local changes to chromatin structure via chromatin remodelling and epigenetic modification. The biophysical properties of the genome may therefore vary significantly over time, from region to region, and from cell to cell.

Semifleixble polymer models are frequently used to decipher the spatial and temporal aspects of chromosome organisation. Such models allow for parameter estimation from experimental observations (Bystricky et al., 2004, Ding et al., 2006, Koszul et al., 2008, Arbona et al., 2017), and so provide a concise quantification of the state of the system in terms of meaningful biophysical parameters, such as the compaction factor and bending-modulus. Simulation studies using appropriately parameterised models may also provide novel insights, and allow for predictions without confounding pleiotropic effects (Penfold et al., 2012), thus guiding future studies.

Most semifleixble polymer models do not explicitly consider the spatial non-stationarity of chromosomes and chromatin. Furthermore, recent advances in chromosome conformation capture (3C)-based allow chromosome organisation to be (indirectly) measured in single cells (Belton et al., 2012, Nagano et al., 2013, 2016). The increasing availability of ensembles of trajectories sampled from potentially heterogeneous populations of cells means it is of interest to develop polymer statistic models that can capture both the spatial nonstationarity of the biophysical parameters, and the statistical relationships that exist within the population. Here we outline a statistical framework for non-stationary semiflexible polymers, and demonstrate how inference can be performed using ensembles of trajectories. For cells belonging to a homogenous population where the biophysical parameters are approximately identical in all cells, a (transformed) Gaussian process prior is assigned to the bending-modulus, and Markov chain Monte Carlo (MCMC) used to infer the posterior distribution of free parameters. For heterogeneous populations of cells, a transformed hierarchical GP (HGP) prior is assigned to the biophysical parameters, which naturally captures the statistical dependency of the parameters that exist across the population. Simulation studies demonstrate the accuracy of the model for homogenous and heterogeneous populations, while applications to yeast chromosome data demonstrates an improved ability to recapitulate trajectories of held out loci compared to related stationary models.

Code used for these analyses, and a graphical user interface (GUI) for simulating non-stationary semifleixble polymer trajectories, have been implemented in MATLAB, and are available to download from: https://github.com/cap76/3MC.

## 1 Semiflexible Polymer Statistics

Chromosomes can be represented via a one dimensional co-ordinate system, *x* ∈ [0,*l*_*c*_], which denotes position along the chromosome, and *l*_*c*_ represents the contour-length. Experimental observations typically yield direct or indirect spatial measurements for a discrete set of (often uniformly incremented) loci, **x** = {**x**_*i*_: **x**_*i*_ ∈ *x*, ***i*** ∈ ℕ, ***i*** ≤ *N*}, with corresponding positions 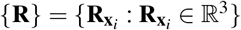, connected via *N* − 1 linking vectors which are usually assumed to be of identical magnitude, 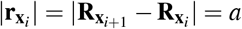. In the discrete worm-like chain (WLC) formulation, the trajectory {**R**} is assigned the following bending energy:

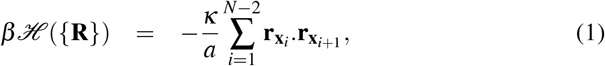

where **κ** denotes the bending-modulus. The probability of a particular configuration may be calculated by awarding each microstate a Boltzmann weighting:

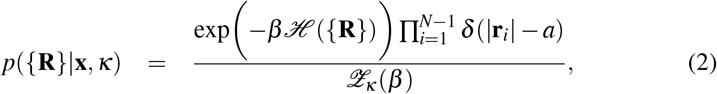

where 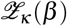 denotes the normalising partition function, and the product of delta-functions ensures inextensibility of the chain.

### 1.1 Non-stationary semiflexible polymers

A heterogenous bending-modulus may be introduced within semiflexible polymer models by a trivial change to Equation (1), to yield:

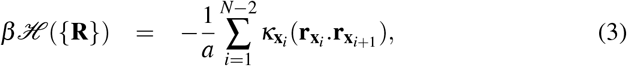

where 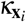 represents a local bending-modulus between links *i* and *i* + 1. Substituting Equation (3) for (1) into Equation (2) increases the parameter space from 3 to (3 + *N* − 2). Since *N* is typically large, constraints must be placed on the bending-modulus for inference to be practical. Here we assume 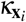 to vary smoothly along the contour-length of the chromosome, and therefore be drawn *a priori* from an appropriate transformation of a Gaussian process:

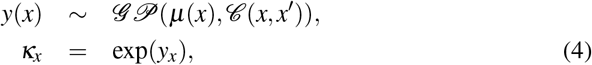

where 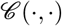 denotes the covariance function. Note that we must take an appropriate transformation of the GP to ensure a positive bending-modulus for all loci. The posterior density over free parameters may be calculated using Bayes’s rule:

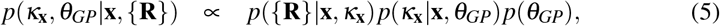

where θ_*GP*_ represents the hyperparameters of the Gaussian process prior and *p*({**R**}|**x, κ**) is given by Equation (2). For confined non-statioanry WLCs, the partition function is analytically intractable, but may be estimated using MCMC approaches (Penfold et al., 2012). However, if the chromosome is unconfined with no other external potentials, this function is analytically tractable (Kleinert, 2009), with the following solution:

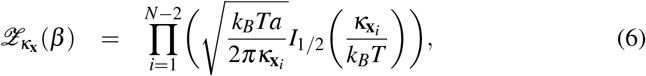

where *I*_1/2_(·) represents a modified Bessel function of the first kind.

### 1.2 Statistical models for ensembles of trajectories with identical bending-modulus profiles

Next we consider a population of cells with corresponding trajectories, {**R**}^(1,…,*M*)^ = {{**R**}^(1)^,…, {**R**}^(*M*)^}, drawn from a homogenous population i.e., with approximately identical bending-modulus profiles. Here we note that the individual trajectories are conditionally independent of one another given the biophysical parameters, and the posterior density over free parameters is given by:

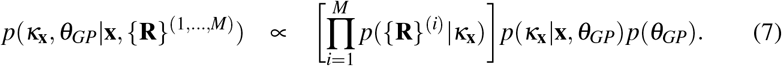

### 1.3 Statistical models for ensembles of trajectories with correlated bending-modulus profiles

An obvious extension to the previous model is for an ensemble of cells draw from a heterogenous population i.e., where the bending modulus profile is non-identical between cells, but may be statistically related. Within a developmental process, for example, the parameters for cells at any given developmental stage may be closely related to the parameters for cells of the preceding stage. For the case in which there exists *M* cells whose parameters are related to one another in a Markovian manner, we can assume the following hierarchical model for the latent bending-modulus profile:

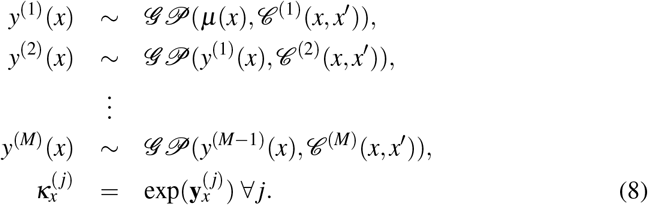

This represents an example of a hierarchical Gaussian process (HGP, Hensman et al. 2013), and we can thus specify a HGP prior over the bending-moduli to arrive at the posterior distribution:

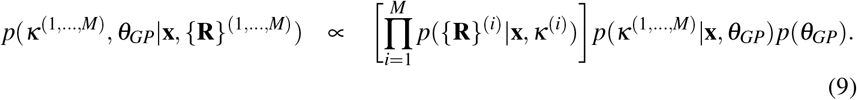

Alternative statistical relationships can be induced between the parameters of individual cells using HGPs, with a notable example being the case where parameters for each cell represent a perturbation of a single latent process, *h*(*x*):

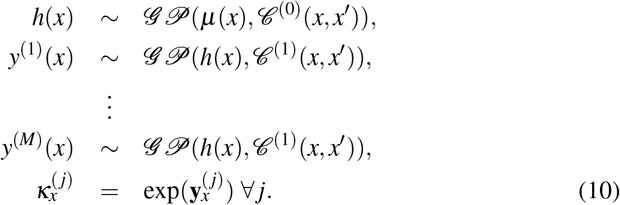

Another useful case exists for heterogeneous populations where, instead of *M* cells within the population, there are *K* statistically related subgroups, where the biophysical parameters within a subgroup are approximately identical. We therefore have an ensemble of trajectories, 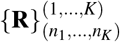, where 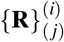 denotes the *j*th trajectory of *n*_*i*_ in subgroup *i*. Using Bayes’s rule we arrive at the following posterior distribution:

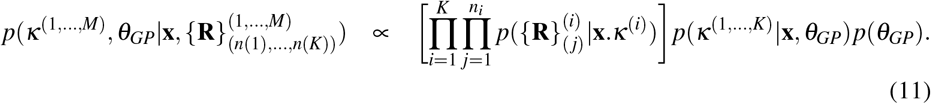

### 1.4 MCMC Algorithm

The posterior densities outlined in previous sections can be efficiently sampled using an MCMC procedure. An initial latent trajectory for the bending-modulus profile is generated as a independent sample from an appropriate Gaussian process prior. All remaining parameters may be set as an i.i.d. samples from the prior and the (unnormalised) posterior density of the model calculated.

A trial bending-modulus trajectory, **κ**\*, may be generated as a sample from a Gaussian process with appropriately chosen control variables (Lawrence et al., 2009). Specifically, *P* control variables are drawn without replacement from the full set of loci **x**, to yield the control input set **x**\* ⊂ **x** and associated control variables **κ**\*. In order to allow for efficient exploration of the posterior process, the cardinality of the control input set, denoted |**x**|, may be assumed to be Binomially distributed with tuneable parameter *p*. An independent sample from a GP is thus generated for the entire set **x** using the *M* control variables as:

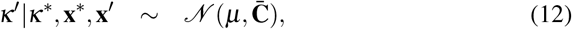

where **x**′ = **x**\**x**\* denotes the complementary set of loci and:

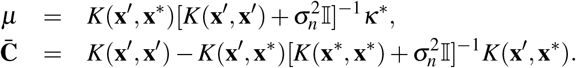

The trial bending-modulus may then be accepted with probability min(1, *A*), where

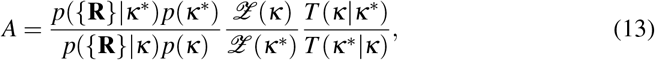

where:

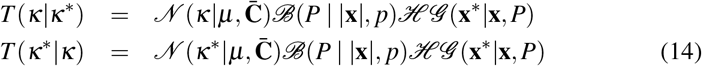

where the second and third terms will cancel in Equation (10) to yield a ratio of Gaussian densities.

## 2 Inference for non-stationary semiflexible polymers

An extension to the 3MC GUI (Penfold et al., 2012) was used to simulate chromosome trajectories as semifleixble polymers with nonstationary bending-modulus profiles. Simulated chromosomes were assumed to correspond to untethered, unconfined WLCs with bending-modulus at locus **x**_*i*_ given by:

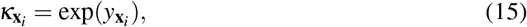

where 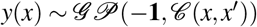 denotes a Gaussian process with constant mean function *μ* (*x*) = −1, and squared-exponential covariance function 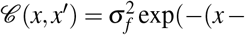, with generative hyperparameters set to ln *l* = 0.1 and ln σ_*p*_ = 0.3.

A number of sample trajectories are shown in Figure 1(a). We first attempted to recover the bending-modulus profile using MCMC as the number of observed trajectories, *N*_*t*_, was increased (*N*_*t*_ ∈ {1,2,4,8,16}). A total of 300,000 samples were generated, with the first 50, 000 discarded for burn-in. The number of control variables used to update the latent GP was assumed to be Binomially distributed, with tuneable probability parameter, *p*, so as to allow optimal acceptance rates. In Figure 1(b) we indicate the inferred posterior bending-modulus profile and 99% confidence intervals (CI) versus the true generating function (sold red line), when hyperparameters of the GP prior were fixed to the correct generative values, ln *l* = 0.1 and ln σ_*p*_ = 0.3. In Figure 1(c) we indicate the inferred bending-modulus profile (with 99% CIs) versus the true generating function (sold red line), when hyperparameters of the Gaussian process prior were incorrectly set as ln *l* = 0.05 and ln σ_*p*_ = 0.05. Similarly, in Figure 1(d) we indicate the inferred bending-modulus profile when the Gaussian process hyperparmeters were also tuned within the MCMC sampling scheme.

**Figure 1:**
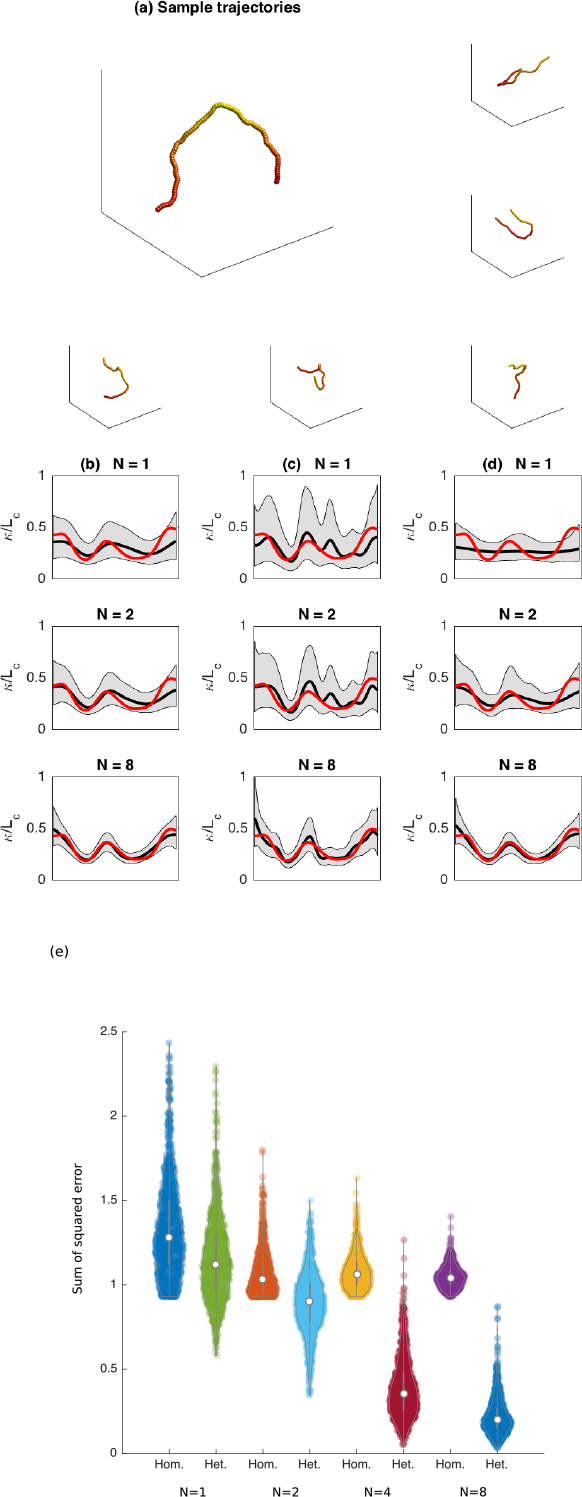
(a) Example trajectories for a semiflexible polymer with non-stationary bending modulus, 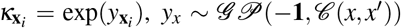, where 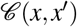 is a squared-exponential covariance function with length scale, log *l* = 0.1, and process-variance, log σ_*f*_ = 0.3. (b) Inferred posterior distribution for the bending-modulus profile (and 95% confidence interval) as the number of observed independent trajectories increases (*N* = 1, 2, 8), when Gaussian process prior hyperparameters were set to the values used for the generating model (log *l* = 0.1, log σ_*f*_ = 0.3), with the true bending-modulus profile is shown in red. (b) Inferred posterior distribution for the bending-modulus profile (and 95% confidence interval) as the number of observed independent trajectories increases (*N* = 1,2,8), when the Gaussian process prior hyperparameters were set to the incorrect values (log *l* = 0.05, log σ_*f*_ = 0.5). (c) Inferred posterior distribution for the bending-modulus profile as the number of observed independent trajectories increases, when Gaussian process prior hyperparameters were tuned with the inference procedure. (d) Even for relatively few observations *N* = 2, the model appears to capture the underlying bending-modulus profile, with reduced sum squared errors compared to wormlike chain models with stationary bending moduli.

Even when relatively few trajectories were observed, the true generating function tended to lie within the estimated 99% confidence intervals of the posterior process, even when the GP hyper-parameters were fixed to incorrect values. As the numbers of observations increased there appeared to be a closer match to the true generating function, with noticeably tighter error-bars. In Figure 1(e) we indicate the distribution of the sum of squared error (SSE) across the observed loci for a heterogeneous WLC with tuned GP hyperparameters compared to MCMC sampling of a homogenous WLC. The results suggest that, in general, the heterogeneous WLC model performs better than the homogenous WLC with no obvious overfitting, with decreased average SSE even for single observations. Thus, heterogeneous biophysical parameters can be recovered from ensembles of trajectories from homogenous populations of cells.

### 2.0.1 Non-stationarity in yeast chromosomes

We next used our approach to infer the bending-modulus profile for *Saccharomyces* cerevisiae chromosome I during interphase, using trajectories of Duan et al. (2010). Here chromosome I was represented by 439 approximately uniformly incremented loci. As with previous analyses, we generated 300, 000 samples in the MCMC chain, with the first 50,000 discarded for burn-in, with the number of control variables tuned to ensure optimal acceptance rates. The inference suggests that whilst chromosome I was reasonably homogenous over relatively large regions, a number of loci were noted that appeared distinctly more rigid. Whilst the true bending-modulus for chromosome I could not be experimentally verifiable, we could assess how well simulated trajectories for a subset of loci resembled those of the observed data. In particular we generated samples form the conditional distribution, 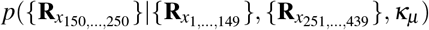, where **κ**_*μ*_ denotes the empirical mean of the bending modulus or bending modulus profile using either a stationary or non-stationary semiflexible polymer model. In Figure 2(a,b) we indicate simulated trajectories for loci 150 to 250 conditional on all other loci and the estimated bending modulus, and in Figure 2(c) indicate the distance between simulated trajectories and the underlying trajectory for those loci, which shows that heterogeneous wormlike chains more closely mimic the underlying trajectories than homogenous wormlike chains, with reduced SSE.

**Figure 2:**
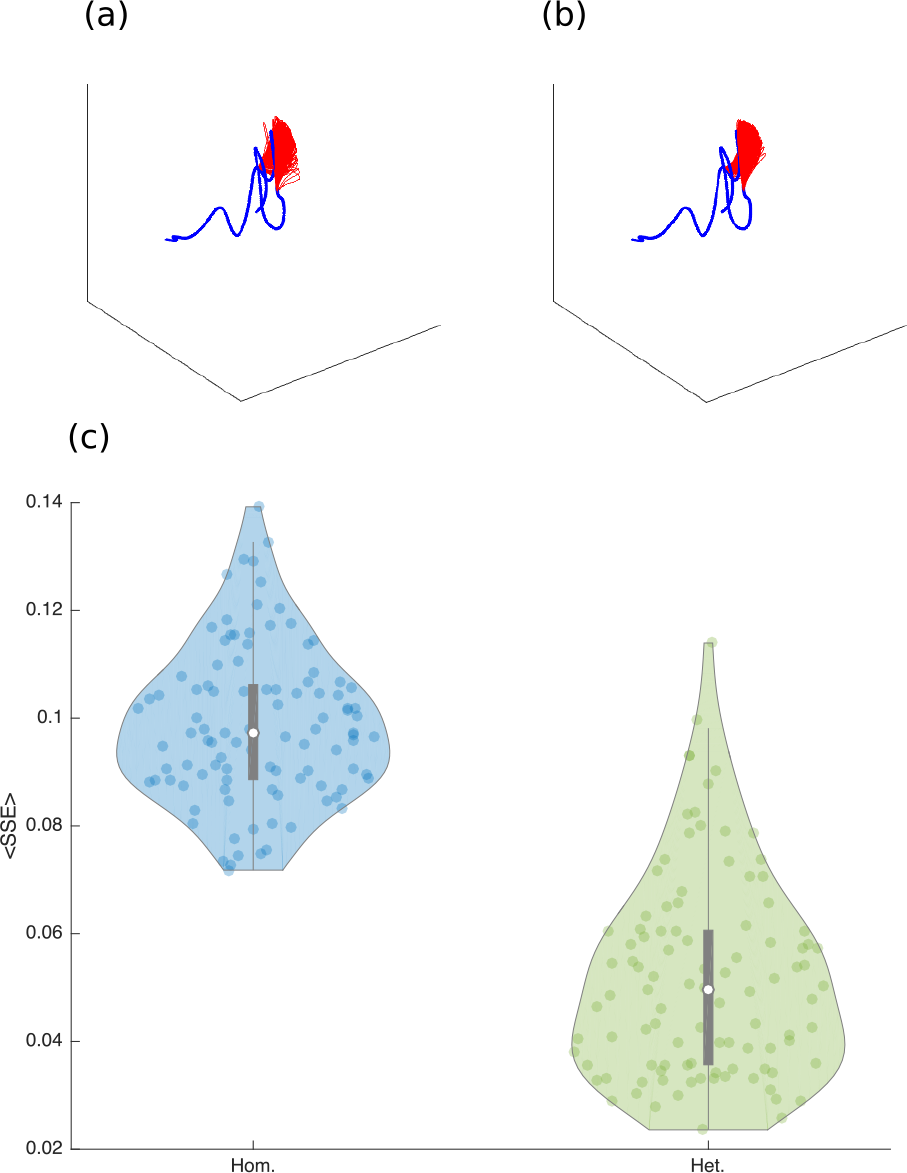
Trajectories for yeast chromosome I (solid blue line) from Duan et al. (2010), along with independently sampled trajectories for loci 150 to 250 (red lines) using the mean bending modulus of a homogenous WLC (a) and mean bending modulus profile of a heterogeneous WLC (b). Simulating trajectories using a heterogeneous WLC model results in reduced SSE (c).

## 3 Discussion

Here we have outlined a method for inferring parameters in non-stationary semiflexible polymers from ensembles of trajectories. For homogenous ensembles we do so by assigning the bending-modulus of the WLC a GP prior, whilst for heterogeneous ensembles, and particularly those comprised of a number of relatively homogenous subgroups, we do so by assigning a hierarchical GP prior, which naturally allows us to capture the statistical dependencies that exists across the groups.

Within this paper we have assumed simple semiflexible models, specifically the case where the chromosome is untethered and far from boundaries. For this case the partition function in the polymer statistical model is tractable even for nonstationary bending-modulus profiles, allowing for feasible inference. However, more interesting cases occur when chromosomes are tethered, or where they lie close to boundaries or domains. In these situations it will be necessary to infer these constants, making the model doubly intractable. Although these constants can be easily simulated via MCMC approaches, the need to do so makes inference less accessible, and approximate methods will be necessary to make these methods applicable. One such approach could be to utilise machine learning approaches, such as independent Gaussian processes, to infer some constants from a more limited number of simulation based studies.

Another key limitation to this work is that inference requires the observation of chromosome trajectories. However, current approaches for investigating chromosome architecture, such as those methods derived from chromosome conformation capture, instead provide indirect measures of chromosome trajectories. In order to be useful, and be more than a illustrative toy model, future development will need to be able to perform inference directly with these indirect measures. Due to the rapid advancement of the field, this is currently beyond my expertise, as much of the work for this paper was started nearly a decade ago. My hope is that the accompanying code of this paper may be useful to others pursuing this line of research, even if only as an illustrative example of a dead end.

## Acknowledgement

Aspects of this work was instigated during my PhD studies, and I’m therefore grateful to my supervisors Professors Alastair S. H. Goldman and Neil D. Lawrence, and the funding I received from the BBSRC. Further developments were made during my time at Warwick, and I’m grateful to Professor David Wild, and funding from the BBSRC (BB/F005806/1) and EPSRC (EP/G021163/1).

